# An efficient field and laboratory workflow for plant phylotranscriptomic projects^1^

**DOI:** 10.1101/079582

**Authors:** Ya Yang, Michael J. Moore, Samuel F. Brockington, Alfonso Timoneda-Monfort, Tao Feng, Hannah E. Marx, Joseph F. Walker, Stephen A. Smith

**Affiliations:** Department of Ecology & Evolutionary Biology, University of Michigan, Ann Arbor. 830 North University Avenue, Ann Arbor, Michigan 48109 USA; Department of Biology, Oberlin College, 119 Woodland St, Oberlin, OH 44074-1097; Department of Plant Sciences, University of Cambridge, Cambridge CB2 3EA, UK.; Department of Ecology & Evolutionary Biology, University of Arizona, Tucson, AZ 85721, USA; Current address: Department of Plant Biology, University of Minnesota-Twin Cities. 1445 Gortner Avenue, St. Paul, MN 55108 Email addresses: YY: MJM: Michael. SFB: AT: TF: HEM: JFW: SAS

**Author notes:** Manuscript received__________; revision accepted__________.

**Keywords:** Caryophyllales, cryogenic field sampling, phylotranscriptomics, phylogenomics, RNA

## Abstract

- *Premise of the study:* We describe a field and lab workflow developed for plant phylotranscriptomic projects, involving field collected cryogenic tissues, RNA extraction and quality control, and library preparation. We also make recommendations for sample curation.
- *Methods and Results:* 216 frozen tissue samples of Caryophyllales and other angiosperm taxa were collected from the field or botanical gardens and were subjected to RNA extraction, stranded mRNA library preparation and sequencing on Illumina HiSeq platforms. These include difficult mucilaginous tissues such as those of Cactaceae and Droseraceae.
- *Conclusions:* Our workflow is not only cost effective (~$270 per sample, as of August 2016, from tissue to reads) and time efficient (~5 hours a sample including all lab work and sample curation), but has proven robust for extraction of difficult samples such as tissues containing high levels of secondary compounds.

## INTRODUCTION

Using *de novo*-assembled transcriptomes to investigate phylogenetic relationships and gene family evolution in non-model plants, or phylotranscriptomics, has gained popularity in recent years due to decreases in cost and improvements in analysis pipelines (Wickett et al., 2014; Edger et al., 2015; Li et al., 2015; Yang et al., 2015; McKain et al., 2016). It is often possible to recover half or more of the total genes in a genome using *de novo*-assembled transcriptome data (Yang and Smith, 2013). Given that a typical plant genome has around 30,000 genes, this would be approximately 15,000 genes per transcriptome, approximately 5,000 of which are shared among most data sets (Yang et al., 2015), with the rest being tissue- and/or taxon-specific. Together they provide enormously rich data for both phylogenetic reconstruction and for investigating gene family evolution that underlies lineage-specific adaptations.

Previous literature on phylotranscriptomic methods has focused on RNA extraction (Johnson et al., 2012; Yockteng et al., 2013; Jordon-Thaden et al., 2015) and data analyses (Yang and Smith, 2013; Yang and Smith, 2014). However, as phylotranscriptomic studies expand to non-model systems that often require field sampling, the logistics of obtaining fresh tissues becomes the limiting factor. In addition, taxa with high levels of secondary compounds, like cacti, are still a challenge for RNA extraction (Jordon-Thaden et al., 2015). As the emerging field of phylotranscriptomics moves forward, the issues of long-term preservation and curation of cryogenic genetic materials will be of the utmost importance for labs seeking to pursue these studies.

From the years 2012 to 2015, we conducted field expeditions to remote localities in both the southwest United States and northern Mexico to support NSF-funded projects on the evolution of Caryophyllales and gypsum endemic plants. Together with samples from living collections, we generated a transcriptomes data set of 200 species of plants (Appendix 1). During the process we have developed an optimized workflow, which is described below. In addition, we discuss alternative procedures that we tested, as well as considerations for project planning.

## METHODS AND RESULTS

### Taxon sampling

The Caryophyllales phylotranscriptomics project emphasized a combination of broad taxon sampling across the order and in-depth sampling of lineages with key evolutionary transitions. These key transitions include the gain and loss of plant carnivory, the gain and loss of betalain pigmentation, transitions to saline, dry, alpine habitats and/or specialized soil types, and transitions to C_4_ and CAM photosynthesis. We planned the field collections in conjunction with a gypsum endemic project, which overlapped partly with the Caryophyllales project in both taxa and geographic region. Of the transcriptomes we have generated for the Caryophyllales phylotranscriptomic project, half were collected from the field, the remaining half from living collections (Appendix 1), with the remaining samples from publicly available transcriptomic and genomic databases such as Phytozome (Goodstein et al., 2012), NCBI Sequence Read Archive (SRA), and the 1000 Plants Initiative (1KP; Matasci et al., 2014).

### Field collection

We timed our field trips to coincide with the beginning of flowering season as much as possible to optimize the chance of obtaining young flower and leaf buds. Our experience has been that mature vegetative tissue is more difficult to work with due to its low concentration of nuclear RNA (Johnson et al., 2012), high level of chloroplast RNA (as they have become specialized photosynthetic apparatus), and high concentration of secondary compounds compared to vigorously growing tissues. It is also important to emphasize that field conditions are more difficult than greenhouse conditions to control. While this may impose limitations for researchers wishing to study differential gene expression, this is not problematic for phylotranscriptomic studies such as ours.

Compared to tissue preservation using an RNA stabilization solution (such as RNA*later*, ThermoFisher Scientific), tissue frozen in the field allows for both DNA and RNA sequencing as well as biochemical analyses, as demonstrated by our ability to use these samples for the characterization of betalain and anthocyanin pigmentation, and hence this was our primary (and recommended) means of collection. For all individuals frozen in liquid nitrogen, we also collected silica-preserved tissue from the same individual as a DNA backup, as well as herbarium specimens whenever possible. Because DNA may degrade relatively quickly for some groups in silica (e.g. Onagraceae), it is important to remove silica from the leaves once dried and place them in a −20°C freezer for long-term storage (Neubig *et al.* 2014).

### RNA extraction, quality control, and DNase digestion

We tested five alternative RNA extraction protocols. These include TRIzol option 1 from Jordon-Thaden et al. (2015), the Aurum^TM^ Total RNA Mini Kit (BIO-RAD) following the manufacturer’s protocol, the QIAGEN RNeasy Mini Kit (QIAGEN) following the manufacturer’s protocol, the PureLink protocol (Ambion; see protocol in Appendix 3; Yockteng et al., 2013) and the hot acid phenol-LiCl-RNeasy Mini Kit protocol [see protocol in Appendix 4, modified from Protocol 12 of Johnson et al. (2012)]. We had approximately 10-30% success rate with BIO-RAD, QiaGen and TRIzol protocols, whereas the PureLink kit had close to 100% success rate and only failed when the sample itself was degraded or highly mucilaginous. For those tissues that are highly mucilaginous like cacti, the hot acid phenol-LiCl-RNeasy Mini Kit protocol was specifically developed to treat these materials. Following RNA extraction, removal of DNA was carried out following Jordon-Thaden et al. (2015) with minor modifications (Appendix 5)

For quality control of RNA, we used agarose gel for an initial assessment. If positive, we followed Fig. 2 of Jordon-Thaden et al. (2015) for evaluating integrity of RNA on a 2100 Bioanalyzer (Agilent). When Bioanalyzer is not available, an alternative method using a NanoDrop Spectrophotometer (Thermo Scientific) was used for an initial assessment of RNA extraction. Positive samples were quantified on a Qubit fluorometer (ThermoFisher Scientific) and a Fragment Analyzer (Advanced Analytical Technologies). When RNA extraction fails, it is often due to either pellet loss (resulting in a completely empty gel with no DNA or RNA trace) or degradation (which shows up as smeared ribosomal RNA bands). RNA degradation can happen during collection, shipping, or in a sub-optimal extraction, as for example with too much starting tissue. For difficult tissues that are mucilaginous, we reduced the amount of starting tissue by half.

### Library preparation

We tested four different library preparation protocols. In 2012 we started with Illumina TruSeq v2, with and without additional strand specific steps [see Supplementary Methods in Yang et al., (2015)]. In 2013 we began using the Yang et al., p.8 newly released TruSeq Stranded mRNA Library Prep Kit (“the Illumina kit”, Illumina), which was more streamlined and produced much higher strand specificity than the previous stranded protocol. In 2014 we switched to the KAPA Stranded mRNA-Seq kit (“the KAPA kit”, KAPA Biosystems; Appendix 6), which is considerably cheaper than the Illumina kit with indistinguishable results in terms of both success rate and strand specificity. The KAPA kit is also more streamlined with less bead washing steps and roughly 15% shorter handling time. The cost is about $30 per sample for the KAPA kit itself plus approximately $20 per sample for consumables (magnetic beads, tips, tubes, and additional chemicals; we used leftover adaptors from the Illumina kit, which lasted through more than 150 additional libraries from one 48-sample Illumina kit).

### Sample curation

We store all RNAs in a −80°C freezer on standard storage racks. Ideally, they would be stored long-term in liquid nitrogen vapor freezers. To prevent freeze/thaw of sensitive samples, we placed samples into labeled freezer boxes and recorded the sample locations in a database that is properly backed up (Appendix 7).

## CONCLUSIONS

We have developed an effective phylotranscriptomics workflow involving collecting tissues cryogenically in the field, RNA extraction of diverse taxa with close to 100% success rate, library preparation optimized for 125 to 150 bp paired-end, stranded mRNA-seq on the HiSeq platforms, and sample storage and curation. Future efforts should focus on streamline the workflow given specific lab and field settings and as sequencing technologies continue to evolve. In addition, future efforts should also be made collaborating with major tissue and seed banks such as the Millennium Seed Bank at Kew and the Smithsonian Global Genome Initiative (Gostel et al. 2016).

## Acknowledgments

We thank H. Flores Olvera, H. Ochoterena, N. Douglas, A. Clifford, S. Lavergne, T. Stoughton, N. Jensen, W. Judd, U. Eggli, G. Kadereit, R. Puente, L. Majure, D. Warmington, S. Pedersen, K. Thiele for assisting with obtaining plant materials; Bureau of Land Management, US Forest Service, California State Parks, Missouri Botanical Garden, Rancho Santa Ana Botanic Garden, Desert Botanical Garden, The Kampong of the National Tropical Botanical Garden, Sukkulenten-Sammlung Zürich, Cairns Botanic Gardens, Botanischen Gartens--Technische Universität Dresden, Millennium Seed Bank, and Booderee National Park for granting us access to their plant materials; and M.R.M. Marchán-Rivadeneira, L. Cortés Ortiz, S. Ahluwalia, J. Olivieri, V. S. Mandala, R. Mostow, M. Croley, L. Leatherman, R. Cronn, M. Parks, T. Jennings, and I. Jordon-Thaden for help with developing lab protocols. The molecular work of this study was conducted in the Genomic Diversity Laboratory at the University of Michigan. Support came from University of Michigan, Oberlin College, the National Geographic Society, a Chateaubriand Fellowship, and the National Science Foundation (DGE 1144254, DEB 1054539, DEB 1352907 and DEB 1354048).

## APPENDICES

## APPENDIX 2.

### Field collection with liquid nitrogen: two alternative setups

Setup 1: Collecting along the road or within a short hike by driving to field sites (Prepared by Michael Moore and Ya Yang)

Field collection setup, with the trunk of the field vehicle doubling as storage and as a wind-blocking, sample processing workbench.

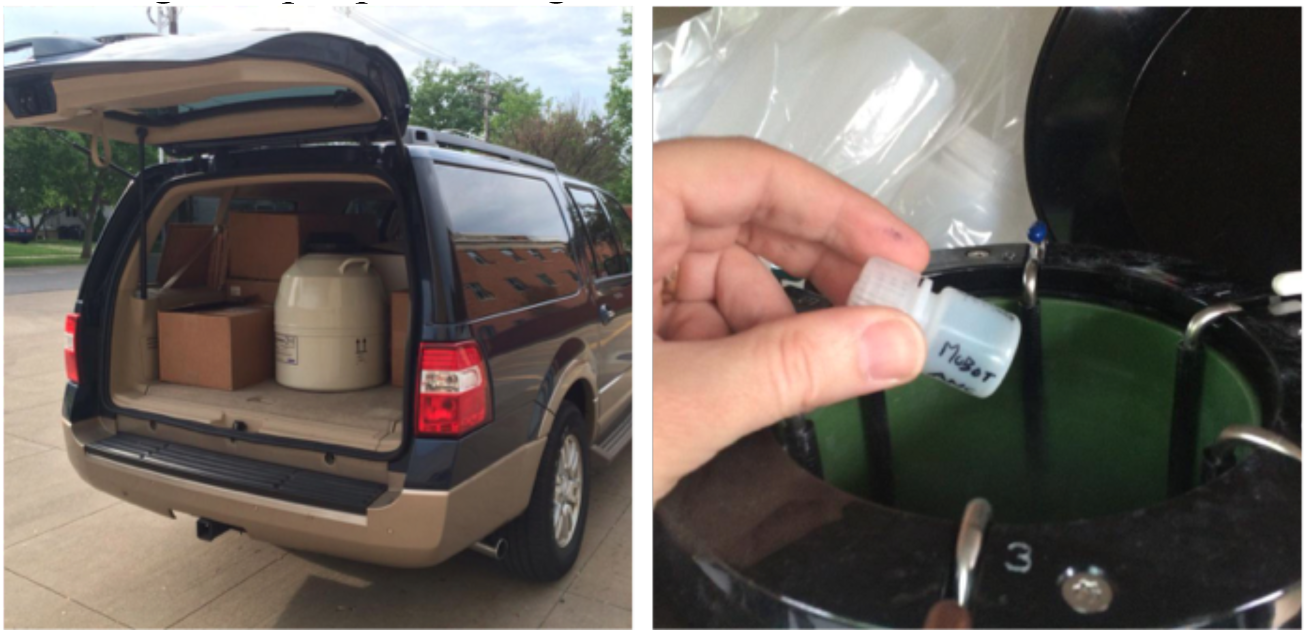

Field supplies:

1. Plant press, straps, cardboard, blotting paper, and newspaper; (optional) field press
2. Coin envelopes for seeds
3. 2" x 3" 2 Mil. clear reclosable bags, 1 per sample
4. GPS unit and maps
5. Black sharpies (blue rub off more easily), pens
6. Field notebook
7. Silica gel in bulk
8. Coffee filters for placing leaf samples in, to be secured using large paper clip and dried in silica gel. Alternatively can use tea bags and small stapler with staples to close tea bags
9. Field guide and keys
10. Hand lens
11. Voucher shipping supplies: shipping tape, strings for tying up specimen into 2-inch bundles with a cardboard on both ends, and cardboard shipping boxes
12. Tools: clippers, Hori-Hori, hammer, scissors
13. Liquid nitrogen tank (shown in figure, MVE Doble 47, Princeton Cryo, Pipersville, PA, USA)
14. Cryo-Gloves, at least mid-arm length
15. Single edge razor blades
16. Long metal tongs (for example VMR 82027-366)
17. 8 mL Nalgene® Boston Round Bottles, High-Density Polyethylene, Narrow Mouth (VWR 16056-988). 2 per sample.

#### Notes

##### Choice of liquid nitrogen containers

There are many options for appropriate liquid nitrogen containers to bring in the field, including nitrogen dewars of varying sizes and dry shippers that possess an absorbent material that leaves a dry interior. There are pros and cons to both styles of containers: dewars often contain larger interiors but care must be taken with the presence of liquid nitrogen, including proper personal protective equipment such as cold gloves and eye protection. Dry shippers often have very small interiors and are not appropriate for large numbers of samples. We recommend the MVE Doble series containers, which are combination dewars/dry shippers that are designed for medium-term sample storage (up to 2 months in appropriate conditions) as well as shipment. The Doble series containers can be filled to the top, and the exterior of the tank will absorb some of the nitrogen but the interior will maintain liquid (albeit at a decreasing volume) for several weeks. We used the Doble 47 container, which has an interior capacity of 47 L. Filled to the top, the tank has stayed reliably cold for over 4 weeks on multiple trips throughout southwestern North America during the summer months, despite repeated jostling on rough unimproved roads. However, these tanks do occupy space, which must be carefully considered when planning a trip.

##### Methods of freezing plant tissue in the field

We have attempted multiple methods of freezing plant tissue in nitrogen in the field, ranging from placing tissue directly into nitrogen-filled containers to placing tissue into bottles and then placing the bottles into nitrogen. Likewise, we have also experimented with leaving tissue-filled bottles in nitrogen for the remaining duration of a field expedition vs. freezing them in nitrogen and then removing them and placing them in dry ice containers for the remaining duration of a field expedition. The former strategy ensures that samples stay appropriately cold with minimal risk of thawing during travel, but not all bottles/containers can withstand being at the temperature of liquid nitrogen for several weeks. The latter strategy obviates this problem, but comes at the cost of having to obtain dry ice at regular intervals (which may be possible in some areas but is often difficult), often every day of the trip, due to the relatively rapid sublimation of dry ice even within a cooler. Because of this, we recommend the former strategy of placing tissue first into bottles and then placing the bottles into nitrogen and leaving them there until returning to the lab. We recommend placing samples in small thick-walled HDPE bottles of 30 mL size or less depending on tissue size; Nalgene manufactures a wide range of such bottles. In practice, 8 mL bottles have been most useful to us given the number of tissues collected; we have successfully accumulated nearly 500 8 mL bottles within a Doble 47 by the end of a 4-week expedition. It is important to note that the caps will come unscrewed for a small proportion of bottles if placed in nitrogen for an extended period; however, our experience has shown that it is possible to minimize this loss to < 1% of bottles if the caps are screwed on as tightly as possible before being placed in nitrogen. For important samples, we take the precaution of putting at least 2 bottles of tissue into nitrogen to ensure that at least one will survive its time in the tank. Finally, it is important to write a sample number on a sheet of paper that is small enough to be easily placed and retrieved (e.g. 1 cm x 1 cm) within the bottle; writing on the outside of a plastic bottle cannot be counted on to survive several weeks in nitrogen.

Tissue sampling itself should proceed quickly, although there is leeway in how much time can elapse between removing a living plant from the soil in the field and freezing the tissue, depending on the goals of sampling. For our project, where transcript expression levels themselves were not a primary consideration, we generally place samples in nitrogen within 60 minutes of removing the plant from the soil or clipping a branch from a large individule, although even longer times have yielded successful, high quality RNA isolations. If longer than 30 minutes is unavoidable, as might be the case if hiking away several kilometers from the field vehicle to a collecting site, it is important to keep the plant in a bag to keep it moist but not let the bag heat up too much by leaving it in the sun. Prior to placing tissue in sample bottles, it is important to break up tissues into pieces small enough that they can be easily retrieved for RNA isolation, especially for succulent tissue.

#### Field procedure

1. Remove plant material sufficient for RNA, DNA, and voucher material and take it back to the vehicle for processing. Choose at least one plant with many flower buds and young leaves, and the rest with mature flowers and fruits for voucher specimens.
2. Label two Nalgene bottles for each sample. Write collection numbers on the bottle in two places each with a Sharpie so that if one number is rubbed off the other one remains. Put young leaves and flower buds from ONE SINGLE PLANT in both bottles. Choose young and vigorously growing tissue and avoid mature tissue if possible. Also avoid fruits and open flowers to avoid additional alleles once pollinated. For succulent tissue or large flower buds cut the tissue into small pieces using a razor blade into paper punch size. Once they are frozen solid it is much harder to cut a piece for extraction. Switch blazes in between individuals.
3. Write the collection number on a small piece of paper and place it in the bottle after placing tissue in the bottle. This helps ensure that it is easy to remove the paper to check the sample ID without removing plant material. Cut the paper instead of tearing it so that it has smooth edges that will not entangle sample tissue fragments.
4. Close the lid of the bottle as tight as possible and place it into the liquid nitrogen tank for the duration of the trip. Bottles will float in the tank, and will bounce against each other on rough roads, which may cause the Sharpie numbers to rub off but the collection number on the piece of paper inside will be the backup. Although the nitrogen never comes in contact with the tissue directly, the leaves freeze very quickly. In earlier iterations of this sampling protocol, we drilled a small hole into the caps of the bottles to allow nitrogen to contact the tissue immediately, but this resulted in no improvement in transcriptome quality and allowed small fragments of tissue to escape the bottle.
5. Cut a piece of coffee filter in half. Put around 1-2 g of healthy leaf material from the same plant into the coffee filter, fold it, and secure it with large paper clip so that the material will not come in contact with silica gel directly; this promotes reusability of silica gel by preventing contamination with small fragments of leaf material. Write the collection number on the outside of the coffee filter. Place the coffee filter pack into a small ziplock bag, write the collection number on the bag and fill the bag with silica gel.
6. Press voucher specimens for each collection. Preferably make duplicates; in general we make 3-5 total duplicates per collection. Record collection date, location, habitat, plant habit, color, and other specimen information. See Gostel et al. (2016) for additional information on vouchers.
7. Check silica gel bags and vouchers each evening. Change newspaper and silica gel if they are saturated with water.
8. Once back in lab, with Cryo-gloves on, use a pair of long metal tongs to retrieve bottles from the liquid nitrogen and place them into labeled freezer boxes for storage (see sample curation protocol) in the −80°C freezer, or into plastic bags for shipping. Do this in the same room as the −80°C freezer, so that the bottles go directly into the −80°C as quick as possible. It is helpful to have small styrofoam coolers with dry ice to place bottles in after retrieving them from nitrogen, to aid in sorting the samples without allowing them to thaw, prior to placing them in the freezer. Write box numbers on the cardboard storage box before placing them on dry ice to pre-cool.

Setup 2: Collecting based on a field station or a local research lab by flying to the field site (Prepared by Hannah Marx)

#### Field supplies

1. In addition to the above supply, also bring a field dryer as described in Blanco et al. (2016).
2. 2 L metal lined coffee thermos for transporting liquid nitrogen in the field
3. Instead of 8 mL bottles, use six 2 mL safe-lock tubes per sample (Eppendorf). RNA can be directly extracted from this tube. Carbide beads can be placed in the tube prior to collection (and frozen with the samples) or just prior to extraction.
4. Instead of a 47 L liquid nitrogen tank, use a smaller dewar (10L) that holds liquid nitrogen for a month, for example the 10 L Cryogenic Container Liquid Nitrogen Tank with Straps and Carry Bag from <u>HardwareFactoryStore.com</u>. Available from: https://www.amazon.com/dp/B00B4YO1QA/ref=pd_lpo_sbs_dp_ss_1?pf_rd_p=1944687522&pf_rd_s=lpo-top-stripe-1&pf_rd_t=201&pf_rd_i=B007XYHY82&pf_rd_m=ATVPDKIKX0DER&pf_rd_r=MPJATD1XXTWW9CCKSWY1

All field supplies except the 10L dewar and liquid nitrogen can fit into one duffle bag (15" x 15" x 30") and checked for air travel to remote localities (see photo below)

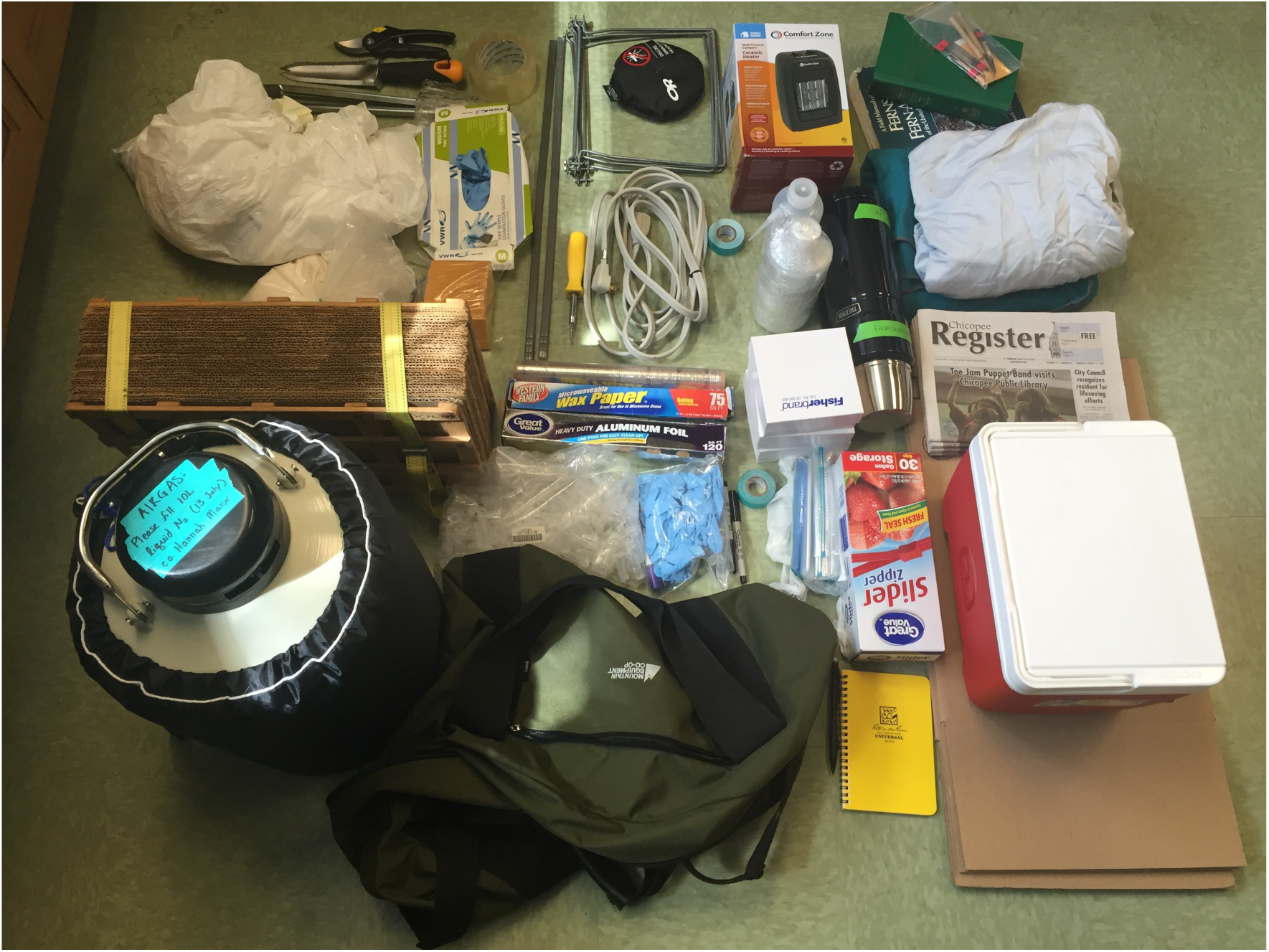

#### Procedure

1. For field sampling, fill the 2 L thermos 3/4 full with liquid nitrogen, and bring this into the field with the cap screwed on half way. Use winter gloves to hold it while hiking.

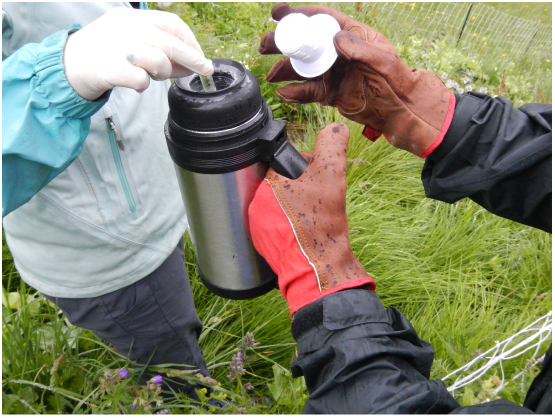
2. For each sample, place about 0.1 g (roughly equal to two hole punches) of mature leaf tissue directly into a 2 mL safe-lock tube, label with the collection number with the black sharpie, and dropped it into the 2 L thermos. Since the tube only stay in the thermos for less than a day we don’t have problem of labels being rubbed off as long as using black sharpies (other colors tend to rub off).
3. Collect 6 replicates for each individual. After finishing, DO NOT SCREW LIDS COMPLETELY. THE LIQUID NITROGEN NEEDS TO VENT TO PREVENT PRESSURE BUILD UP. Collect silica-preserved samples and vouchers as detailed in Gostel et al. (2016).
4. At the end of the day, transfer and organize sample tubes into freezer boxes. Store freezer boxes temporarily on dry ice (if still in the field) or in a −80°C freezer (if near a lab).
5. At the end of field trip, ship three replicates for each individual back on dry ice for extraction, and save the remaining three as backup, usually stored in a lab at a research station near the location where they were sampled.

## APPENDIX 3.

### RNA Extraction using the PureLink reagent

(Approximately 3 to 4 hours. Prepared by Ya Yang and Michael Moore)

#### Planning/Overview

I usually process 6 samples per day and 2 tubes per sample. This enables me to try different tissue types (flower buds vs. leaf) or different amount of tissue (more vs. less if only vegetative tissue is available). 12 tubes at a time also work the best with our 24-place standard room temperature centrifuge. Extraction involves lots of handling time and little wait, and you need to move quickly to avoid RNA degradation so there is little benefit to extracting more at a time. Free crushed dry ice can usually be obtained in departmental biostores when a shipment arrives. Since the entire procedure is carried out in a fume hood, make sure it is available during for the whole day of extraction.

- Day 1: RNA extraction of 6 samples in 12 tubes.
- Day 2: RNA extraction for another 6 samples in 12 tubes. Proceed to DNase digestion and Bioanalyzer for all 12 samples. Normally at least one of the two tubes per sample will work out and Bioanalyzer takes 12 mRNA samples per run.
- Day 3 and Day 4 (or once having 12 samples passing quality control): library prep. Currently we multiplex 10-11 libraries per lane on the HiSeq2500V4 platform.

#### General considerations for working with RNA

- RNA is a less stable molecule than DNA and is prone to degradation. RNases are also abundant within plant tissue and readily degrade RNA. All reagents, containers, and tips used for RNA-related work should be RNase-free. Clean the work surface and pipettes with RNase*Zap* before use. But don’t be paranoid about it. Unless you are being particularly unclean, RNase contamination is not the cause of most failed RNA extractions.
- RNases are not removed by autoclaving so avoid using glassware. Use 15 mL and 50 mL RNase-free centrifuge tubes to aliquot reagents.
- Keep tissue frozen till PureLink is added, which deactivates RNase.
- Regularly replace electrophoresis buffer if RNA is to be visualized in agarose gel, ideally each time before running new samples.
- Always wear disposable gloves and change them frequently
- Avoid freeze and thaw of RNA. Place RNA samples in 4°C if processing the next day or two and store in −80°C if processing later.

#### Tools and equipments

- Access to a fume hood during the entire duration of extraction
- Tweezers with insulated handle and smooth tip (for easy cleaning).
- Tissue homogenizer. We currently use FastPrep-24™ benchtop homogenizer with CoolPrep™ 24X2 mL adapter (MP Biomedicals, Santa Ana, California, USA).
- Tube rack for holding lysing matrix tubes in liquid nitrogen. We currently use the CoolRack® Thermoconductive Tube Racks (BioCision, San Rafael, California, USA) to prepare frozen tissue before homogenization. Certain plastic racks work as well but some will crack. If using a plastic rack, drill a hole at the bottom of each well to allow liquid nitrogen to go through. You may also need to cut a plastic rack short so that it fits into a small styrofoam shipping container.
- Two styrofoam shipping boxes with lids. One is to hold liquid nitrogen and rack and you want a shallow one for easy maneuverability. The other is to hold dry ice.
- Waste beaker
- Liquid nitrogen
- Benchtop liquid nitrogen container (for example, Nalgene® Dewar Flasks, High-Density Polyethylene, Thermo Scientific)
- A room temperature centrifuge
- A refrigerated centrifuge
- A set of designated pipettes for RNA work: P1000, P200, and P20
- Vortexer
- For handling tubes sitting in liquid nitrogen, use a tight fit lightweight insulated glove (for example I use the North Face Powerstretch Glove) underneath latex glove on one hand for picking up tubes, and a latex glove only on the other hand for holding tweezers that has insulated handle

#### Reagents

- Squirt bottle with 70% ethanol
- PureLink® Plant RNA Reagent (Ambion^TM^). Store in 4°C.
- RNase-free water (store in 4°C and aliquot in 50 mL tubes on bench)
- 75% ethanol. Store in 4°C. I usually make 48 mL at a time with 36 mL 200 proof ethanol + 12 mL RNase-free water in a 50 mL centrifuge tube. It is good for 48 extractions.
- 5 M NaCl solution in 50 mL tubes on bench. Use 2.922g NaCl powder for each 10 mL of final volume. Close the lid of a centrifuge tube and invert it back and forth to dissolve. It takes some patience. NaCl is stored in powder.
- Chloroform in non-inflamable cabinet. It is light sensitive and dissolves plastic so only aliquot at use.
- Isopropyl alcohol stored in fire proof cabinet. Aliquot in 50 µL tubes on bench.
- 3 M KOAc, pH 5.2 (optional, for mucilaginous tissue)

#### Consumables

- Kimwipe
- Paper towel
- RNase-free 1.5 mL, 2 mL, and 50 mL tubes
- RNase-free barrier tips (1000uL, 200uL, and 20uL)
- Lysing Matrix A in 2 mL Tube (MP Biomedicals)
- RNase*Zap*® RNase Decontamination Solution (Ambion^TM^)
- Disposable latex gloves

#### Safety

The PureLink reagent contains 2-mercaptoethanol and sodium azide.

- Sodium azide may react with lead and copper plumbing to form explosives. Do not pour down the drain.
- The reagent has a strong odor and causes headache and dizziness when inhaled. Work in the hood and temporarily dispose of tips and tubes in the hood in a ziplock bag. At the end of the day, seal waste bags air tight with double layers of ziplock bags and put in hazardous waste disposal. Otherwise the entire hallway will become stinky!

#### Sample Preparation

1. Fill the benchtop liquid nitrogen container with about 3 L of liquid nitrogen. Obtain ⅓ box of crushed dry ice. Place the Coolrack (or other rack you choose to use) in the shallow styrofoam box, and pour about 1 L of liquid nitrogen into the styrofoam box. Let the rack to chill for a few minutes. Pre-cool the refrigerated centrifuge to 4°C. Check the refrigerated centrifuge to make sure that it has the microtube adapter instead of the plate adapter in it to make sure that the correct adapter is cooled. Switch the adapter if needed and restart the centrifuge.
2. Wipe the workspace with 70% alcohol followed by RNase*Zap*.
3. Write numbers 1-12 on 12 lysing matrices. Tap down the beads. Slightly loosen the caps so that they are easy to unscrew in liquid nitrogen. Place the lysing matrices on rack in liquid nitrogen to let them chill.
4. Gather the following: tweezers with smooth tips, 70% ethanol, RNase*Zap*, Kimwipes, a jar for collecting used tips and tubes, lab notebook, pen, insulated gloves and latex gloves, styrofoam box with dry ice, lab notebook, and laptop. Take the box of dry ice to the freezer room. Take the tissue storage box out of the −80°C freezer and immediately place it on dry ice.
5. Spray the tweezers using 70% ethanol, wipe with a Kimwipe, apply RNase*Zap* and wipe again with another Kimwipe. Twist open the bottle and put lid on side, check the sample number on the bottle and on the paper slip inside and put the slip in the bottle lid on the side. Dip the tweezers in liquid nitrogen to chill. Remove < 0.1 g tissue from the bottle (size of punch hole approximately, I usually don’t weigh them so that I can move quickly to avoid thawing samples). Record tissue types on lab notebook. If unsure about whether a bud is part of an inflorescence or vegetative tissue, Google search the species to find out or ask.
6. Clean and prepare the tweezers by spraying with 70% ethanol and RNase*Zap* as in step 5, and dip in liquid nitrogen to chill before proceeding to the next sample. Add liquid nitrogen to the styrofoam box when it’s low.
7. Tape the cryogenic adapter except 12 (six on each side and make) so that the dry ice won’t fly out while shaking. Transfer < 0.5 g crushed dry ice to the FastPrep adapter. Use small pieces so that it’s easier to balance.
8. Grind frozen tissue in the FastPrep-24 on Cryogenic cycle and 4 m/s for 40 seconds. After finishing, put lysing matrices back in the rack in liquid nitrogen. Tap down the bead on bench while waiting for 5 min required by the FastPrep. Don’t tap too hard since now the tubes are brittle and may crack. Add liquid nitrogen to the styrofoam box if needed. Check leftover dry ice in the adapter. There should be a small amount of fine powder left. If the ground frozen tissue thaws at any point before adding the extraction buffer you will get degraded RNA.
9. Add more dry ice to the adapter and shake for another 40s. Put lysing matrix back on the rack in liquid nitrogen. The tissue should be in very fine powder. If not, repeat for a third round of grinding.

#### RNA extraction

1. Move the foam container with samples in liquid nitrogen and vortexer to the hood and complete all following steps in the hood. Line the waste beaker with a ziplock bag. Move the styrofoam box with samples in liquid nitrogen into the hood. The samples need to be kept frozen until PureLink reagent is added.
2. Take the PureLink reagent out of the 4°C fridge and aliquot 6.3 mL into a 50 mL falcon tube. Tap the frozen tube gently on the counter before opening it so that the beads and most of the powder are at the bottom of the tube instead of stuck to the lid. Add 0.5 mL cold (4°C) PureLink reagent to the frozen, ground plant tissue. Tighten the lid before vortexing the tube until the sample is thoroughly resuspended with no clumps at the bottom of the tube. Put the tube in a clean, room-temperature rack. Return the PureLink reagent back to 4°C fridge.
3. (Optional) Add ⅓ volume KOAc (3 M, pH 5.2) to the lysate. Vortex to mix. This step is used for mucilaginous tissue.
4. Incubate the tube horizontally for 5 minutes at room temperature. While waiting, label 12 1.5 mL tubes with numbers 1-12. Add 0.1 mL of 5 M NaCl to each empty tube.
5. Centrifuge the sample tubes at 12,000 × g for 2 minutes at room temperature.
6. Use 200 µL tips to transfer the supernatant to the new tubes with 5 M NaCl. Do not use 1000 µL tips since liquids within them are more difficult to control. Pipette up and down gently to mix the supernatant with NaCl after transfer.
7. Aliquot 3.6 mL chloroform in two 2 mL tubes. Add 0.3 mL chloroform to each sample. Move quickly so that chloroform does not drip from the pipette tip. Close the lid tight and mix thoroughly by vortex.
8. Centrifuge the sample at 12,000 × g for 10 minutes at 4°C to separate the phases.
9. While waiting, label a new set of 12 tubes with sample ID on top, and date and tube number on side of the tube. Add a volume of isopropyl alcohol equal to that of the aqueous phase (usually 350 to 400 µL) to each empty tube.
10. Transfer the upper, aqueous phase using 200 µL tips to the tubes with isopropyl alcohol added. Make sure not to disturb the middle layer. Mix and let stand at room temperature for 10 minutes. Hold on the waste in the tube and pour them later so that gloves do not get dirty.
11. While waiting, make a 1.5% agarose
12. Centrifuge the sample tubes at 12,000 × g for 10 minutes at 4°C.
13. Decant the supernatant, **taking care not to lose the pellet**. Touch the lip of tube on paper towel to clean up (make sure use a new spot for each tube). Add 1 mL of 75% ethanol to the pellet.
14. Centrifuge at 12,000 × g for 2 minute at room temperature. Decant the supernatant carefully, **taking care not to lose the pellet.** The pellet is even looser than the last round. Touch the lip of tube on designated spot on paper towel to avoid cross-contamination before closing the lid.
15. Briefly centrifuge to collect the residual liquid and remove it with a 20-µL pipette. Leave the tube open to dry for 15–30 min.
16. Add 30 µL RNase-free water to the RNA pellet. Pipet the liquid up and down over the pellet to resuspend the RNA. It is OK if the solution is still cloudy after mixing. It will be cleaned up at the DNase step.
17. Make aliquot of 3 µL for QC. Purified RNA can be kept at 4°C for a day or two, or at - 80°C for long-term storage. Alternatively, proceed to DNase step immediately.
18. Pour waste into waste container. Wash room temperature racks with tap water. Pour waste liquid into the extraction waste collection bottle in the hood. Discard tips and tubes in the sealed ziplock bag to the hazardous waste bucket. Allow leftover dry ice and liquid nitrogen evaporate on the counter and wash the containers and rack sitting in liquid nitrogen the next day.

## APPENDIX 4.

### RNA extraction for mucilage tissue using hot acid phenol-LiCl-RNeasy Mini Kit

[~2 days. Notes and modifications from Protocol 12 in Appendix S1, Johnson et al. (2012). Prepared by Alfonso Timoneda-Monfort and Tao Feng]

#### Equipment

Only listing those that are not required by the default PureLink protocol

- 15 mL RNase-free Falcon tubes (instead of snap cap tube)
- Adapter for 15 mL falcon tubes in the refrigerated centrifuge
- Water bath or dry heating block that holds 15 mL tubes
- Mortar and pestle. Rinse mortar and pestle with water immediately after use, and then rinse with ethanol. We don’t autoclave them because autoclaving will not destroy all RNases. It is ok to have some RNase before extraction buffer is added since plant tissue contains RNases itself.
- We also don’t autoclave the spatulas, we clean them between samples using ethanol, and chilled before touching the powder, otherwise the tissue powder will melt in contact with the metal and stick to it.

#### Reagents

- Saturated acid phenol (pH 4.3)
- Chloroform:Isopropyl alcohol (24:1), RNase free
- Isopropyl alcohol
- 4M LiCl solution
- 70% ethanol made with RNase-free H_2_O, store in 4°C
- Prepare RNA extraction buffer as follows. We did not filter purify them.
Final Concentration:
100 mM Tris pH 9.0
1% SDS
100 mM LiCl
10 mM EDTA
For 100 mL :
10 mL 1M Tris pH 9.0
10 mL 10% SDS
2.5 mL 4M LiCl
2.0 mL 0.5M EDTA pH 8.0
Bring the volume up to 100 mL using RNAse-free water and keep at 4°C

#### Safety

- Phenol could be fatal if inhaled and is rapidly absorbed through skin. Chloroform:isoamyl alcohol solution may cause cancer and damaging the unborn. Work with these compounds must be carried out in a fume hood. Minimize the time tubes are outside the fume hood. Use protective goggles during the whole process and change gloves immediately after any chemical spillage.
- Be careful while adjusting the pH of the Tris solution. Adding HCl will produce toxic vapors of chlorine. Do not inhale; work in the hood.
- Phenol and RNA Extraction Buffer liquid waste should be stored in a separate waste bottle and disposed of separately.

#### Modification to Protocol 12 in Appendix S1, Johnson et al. (2012)

- Starting material: instead of 1 g, we use 0.2 to 0.4 g or even less for Cactaceae.
- In some cases, especially Cactaceae, the pellet is very small and looks clean, and would be lost with the LiCl precipitation. In these cases, skip steps 17–18
- For step 24, elute RNA twice from the column using 65°C RNase-free water instead of 95°C.

## APPENDIX 5.

### DNase digestion

[~1 hour. Modified from the manufacturer's protocol and from Jordon-Thaden et al. (2015). Prepared by Ya Yang]

#### Equipment

In addition to the equipment required for the PureLink RNA extraction protocol, you will need:

- Dry heating block that holds 1.5 µL tubes (preferred), or an incubator
- TURBO DNA-*free*™ Kit (Invitrogen), stored in −20°C freezer
- Agilent 2100 Bioanalyzer and Agilent RNA 6000 Nano Kit (Agilent, Santa Clara, California, USA). Sequencing cores also usually provide Bioanalyzer service.

#### Procedure

1. Take the DNase buffer out of the −20°C freezer to thaw at room temperature. Turn on the dry heater or incubator to preheat to 37°C.
2. Can combine the two tubes per sample to increase yield and diversity of genes (total of ~50 µL). Aliquot 3 µL of RNA sample to a new tube and place in 4°C fridge (if haven’t done so at the end of extraction).
3. Vortex the DNase buffer and spin it down briefly. Add 0.1 volume of 10X Turbo DNase buffer to each tube. For 50 µL of RNA add 5µL buffer.
4. Add 1 µL of DNase from the TURBO DNA-*free* Kit to the RNA. Make sure the 1 µL is indeed pipetted. It can be hard to see. Vortex briefly to mix
5. Incubate at 37°C for 30 minutes. While waiting, label new 1.5 mL storage tubes with the collection number on top, and the date and tube number of extraction on the side.
6. Add vortexed DNase Inactivation Reagent in the TURBO DNA-*free*™ Kit (typically 0.1 volume; 5 µL for 50 µL of starting RNA) and mix by vortex briefly.
7. Incubate at room temperature for 5 minutes, vortex occasionally.
8. Centrifuge at 10,000 x g for 2 min and transfer supernatant to the new, pre-labeled storage tubes, and aliquot 3 µL for bioanalyzer.
9. Use a 1.5% agarose gel to compare the presence of genomic DNA before and after treatment and to test for degradation. It’s OK to use a DNA ladder. Repeat the clean up a second time if a high molecular weight DNA band shows up on the gel. Chloroplast rRNA gives additional bands, and can appear as a smear on a gel but will be distinguishable on Bioanalyzer trace.
10. Place cleaned RNA in 4°C if library prep will be performed in the following day or two. Otherwise store at −80°C.
11. If positive, eun the cleaned RNA on a Bioanalyzer using the Agilent RNA 6000 Nano Kit chips. Mucilaginous tissue can give odd Bioanalyzer traces but library prep might be OK. Take note when mucilaginous tissue that is difficult to pipette occurred during extraction.

## APPENDIX 6.

### Stranded mRNA library preparation

(Approximately 2 days for 12 libraries and 2.5 days for 20 libraries. Prepared by Ya Yang and Michael Moore)

#### Equipment and consumables

In addition to the equipment required for the PureLink RNA extraction protocol, you will need:

- A thermocycler with a dedicated PCR block to store all the programs and ensure access to the machine at all times throughout the duration of the protocol.
- Minicentrifuge for quick spins of both 1.5 mL tubes and PCR strips
- Magnetic-ring stand (96 well). We had one from Ambion (AM10050) but it was quite difficult to use and often resulted in bead loss. We are happy with Agencourt SPRIPlate 96R Ring Magnet Plate (Beckman Coulter, Inc.) that has a stronger magnet.
- AMPure XP beads, 5 mL (Agencourt A63880); larger volumes are available but beads are expensive and only have a shelf life of one year.
- Indexed adapters (we used the leftover adapters from Illumina TruSeq Stranded mRNA library preparation kit. Can also be purchased separately).
- 0.2 mL PCR strips.
- 80% ethanol, 1.6 mL per sample, made fresh for each library prep with RNase-free water.
- 0.01 M Tris-HCl, ph=8. Dilute 100x with RNase-free water from 1M stock solution
- KAPA Stranded mRNA-Seq Kits (KAPA Biosystems).

#### Notes and modifications to the manufacturer's instruction (2015 version)

1. Library prep is carried out in PCR strips. To avoid contamination, do not use multi-channel pipettes and only open one tube at a time. Label PCR strips under the lid to minimize the chance of it being rubbed off.

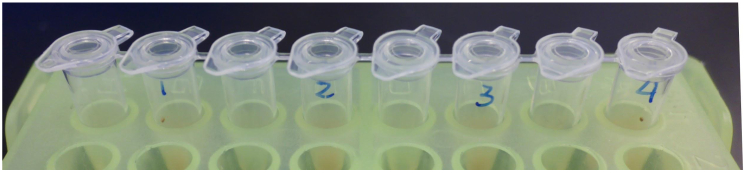
2. Briefly vortex and spin down all reagent tubes taken from the freezer or fridge before opening them.
3. Use the smallest tip (P10 or P2) to pull up the leftover ethanol (usually 1-2 µL) at the end of each bead cleanup while air drying the beads.
4. Since most RNA-seq library prep kits are optimized for differential gene expression studies that uses relatively short read lengths, we modified the protocol to produce larger insertion sizes to accommodate paired-end 125 or 150 bp reads.
  - Fragmentation: 85°C for 6 min.
  - 12 cycles of PCR enrichment
  - Use 0.7x (35 µL) instead of 1x (50 µL) AMPure beads for the final cleanup step after PCR enrichment. This is also more effective in removing adapter contamination.
5. Use 1-2 µL Illumina TruSeq adapters per sample There are other mRNA library kits that claim shorter handling time. Illumina NeoPrep is not working reliably yet as of July 2016, but it looks like a promising future alternative to hand prep.

## APPENDIX 7.

### Sample curation

(Prepared by Ya Yang and Stephen Smith)

This protocol is for curating tissues and RNA samples at a moderate scale (several hundred to a few thousand samples).

#### Considerations on facilities

Freezers fail periodically. Ultra-low freezers should be equipped with a temperature monitor, an alarm system, and should be connected to a backup power generator. Ideally, transfer samples to a liquid nitrogen vapor system for long-term storage.

#### Organizing samples

To fit the 8 mL bottles into standard freezer racks, we use standard storage boxes (2" Cardboard Cryovial Storage Box only, 5 1/4" x 5 1/4" x 1 7/8", Dot Scientific, Inc.) with 16 cell dividers (16 Cell Cardboard Divider, Cell Opening 30.23mm / 1.19", Outside Dimensions 4 7/8" x 4 7/8", Dot Scientific, Inc.). Do not use plastic storage boxes in ultra-low freezers since they get brittle. Do not use tapes since they tend to fall of in freezers.

When organizing bottles into storage boxes upon returning from the field trip, verify the collection number by reading the paper slip inside of the bottle if the number written on outside of the bottle is rubbed off, and record the precise location of each sample in a database. Always sort samples in insulated boxes with ample of fresh dry ice to avoid thawing.

Below shows how we organize sample tubes in cardboard freezer boxes for long-term storage. The collection number can be written on the box cover if needed. Note that identifying information for each box should be clearly indicated on both the cover and the body of the storage box. Cell location ID is recorded as A1, A2 … to D4. All information should be recorded in a database or a spreadsheet that is write-protected and properly backed up.

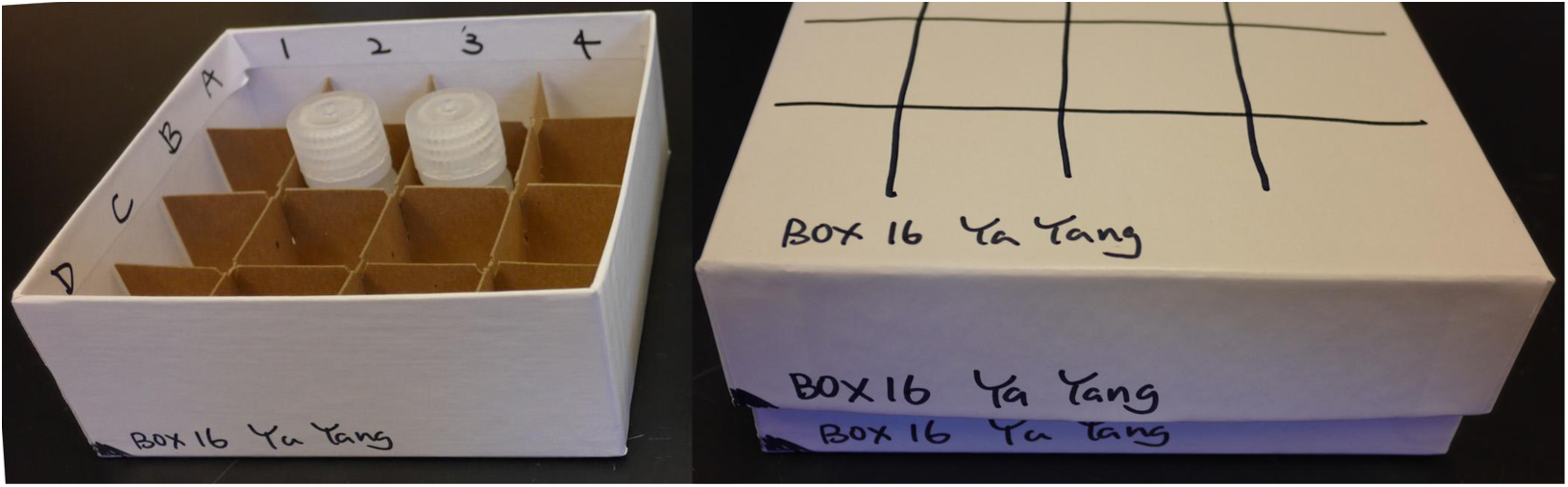

#### Database schema

Below is the database schema we are using to organize sample, extraction and library information, as well as metadata on sequencing reads and assembly. These started as spreadsheets on Google Drive, but we developed an SQL database as the number of samples grew.

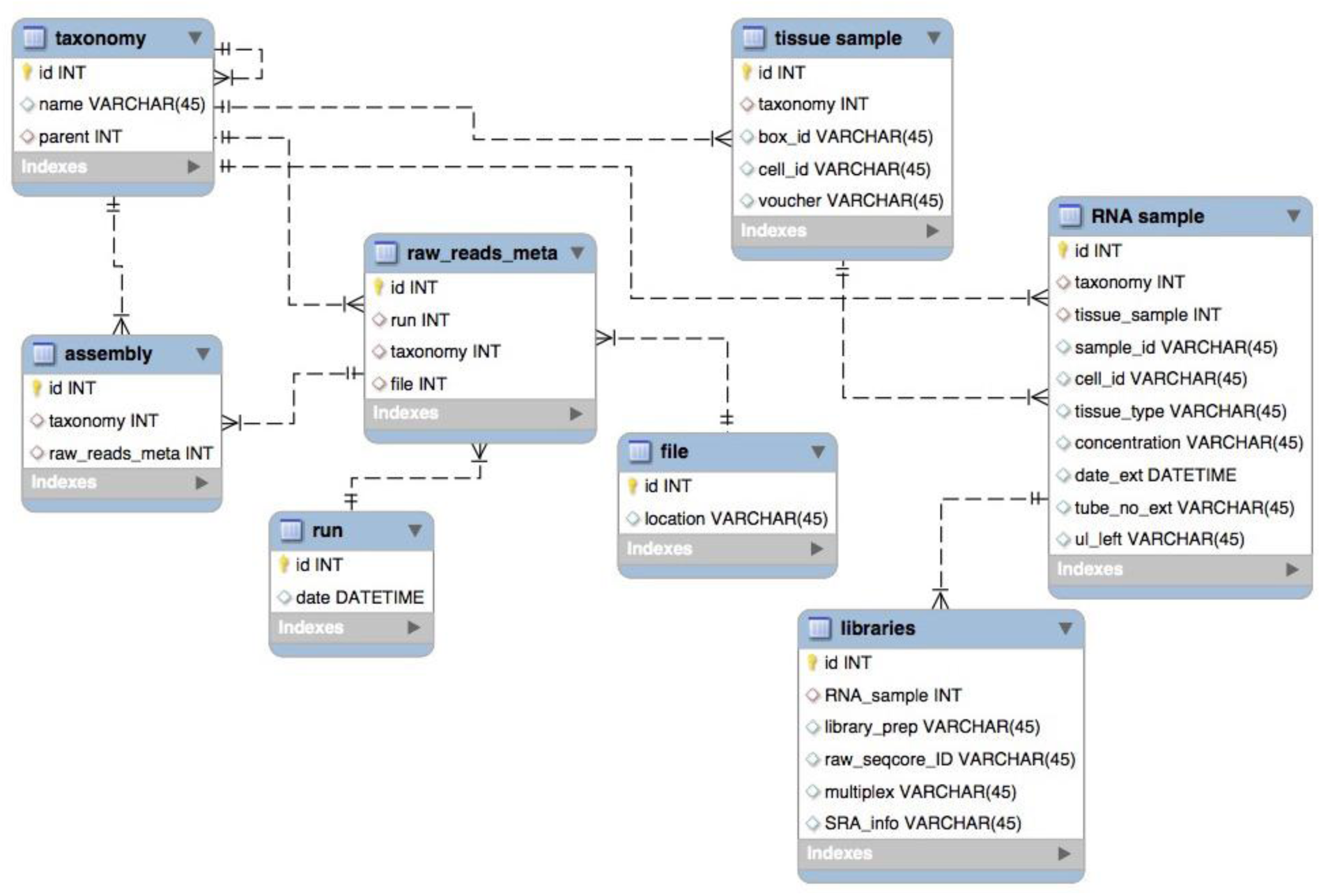

## Literature Cited

Edger, P. P., H. M. Heidel-Fieldischer, M. Bekaert, J. Rota, G. Glöckner, A. E. Platts, D. G. Heckel, et al. 2015. The butterfly plant arms-race escalated by gene and genome duplications. Proceedings of the National Academy of Sciences 112(27): 8362–8366.

Goodstein, D. M., S. Q. Shu, R. Howson, R. Neupane, R. D. Hayes, J. Fazo, T. Mitros, et al. 2012. Phytozome: a comparative platform for green plant genomics. Nucleic Acids Research 40: D1178–D1186.

Gostel, M. R., C. Kelloff, K. Wallick, and V. A. Funk. 2016. A Workflow to preserve genome-quality tissue samples from plants in botanical gardens and arboreta. Applications in Plant Sciences 4: 1600039.

Johnson, M. T. J., E. J. Carpenter, Z. Tian, R. Bruskiewich, J. N. Burris, C. T. Carrigan, M. W. Chase, et al. 2012. Evaluating methods for isolating total RNA and predicting the success of sequencing phylogenetically diverse plant transcriptomes. Plos One 7: e50226.

Jordon-Thaden, I. E., A. S. Chanderbali, M. A. Gitzendanner, and D. E. Soltis. 2015. Modified CTAB and TRIzol protocols improve RNA extraction from chemically complex Embryophyta. Applications in Plant Sciences 3(5): 1400105.

Li, Z., A. E. Baniaga, E. B. Sessa, M. Scascitelli, S. W. Graham, L. H. Rieseberg, and M. S. Barker. 2015. Early genome duplications in conifers and other seed plants. Science Advances 1: e1501084.

Matasci, N., L.-H. Hung, Z. Yan, E. Carpenter, N. Wickett, S. Mirarab, N. Nguyen, et al. 2014. Data access for the 1,000 Plants (1KP) project. GigaScience 3: 17.

Mckain, M. R., H. Tang, J. R. Mcneal, S. Ayyampalayam, J. I. Davis, C. W. Depamphilis, T. J. Givnish, et al. 2016. A phylogenomic assessment of ancient polyploidy and genome evolution across the Poales. Genome Biology and Evolution 8: 1150–1164.

Neubig, K. M., W. M. Whitten, J. R. Abbott, S. Elliott, D. E. Soltis, and P. S. Soltis. 2014. Variables affecting DNA preservation in archival plant specimens. In W. L. Applequist and L. M. Campbell [eds.], DNA banking for the 21st century: Proceedings of the U.S. Workshop on DNA Banking, 81–112. William L. Brown Center, Missouri Botanical Garden, St. Louis, Missouri, USA.

Wickett, N. J., S. Mirarab, N. Nguyen, T. Warnow, E. Carpenter, N. Matasci, S. Ayyampalayam, et al. 2014. Phylotranscriptomic analysis of the origin and early diversification of land plants. Proceedings of the National Academy of Sciences 111: E4859–E4868.

Yang, Y., and S. A. Smith. 2013. Optimizing de novo assembly of short-read RNA-seq data for phylogenomics. BMC Genomics 14: 328.

Yang, Y., and S. A. Smith. 2014. Orthology inference in non-model organisms using transcriptomes and low-coverage genomes: improving accuracy and matrix occupancy for phylogenomics. Molecular Biology and Evolution 31: 3081–3092.

Yang, Y., M. J. Moore, S. F. Brockington, D. E. Soltis, G. K.-S. Wong, E. J. Carpenter, Y. Zhang, et al. 2015. Dissecting molecular evolution in the highly diverse plant clade Caryophyllales using transcriptome sequencing. Molecular Biology and Evolution 32: 2001–2014.

Yockteng, R., A. M. R. Almeida, S. Yee, T. Andre, C. Hill, and C. D. Specht. 2013. A Method for Extracting High-Quality RNA from Diverse Plants for Next-Generation Sequencing and Gene Expression Analyses. Applications in Plant Sciences 1: 1300070.

## Literature Cited

Blanco, M. A., W. M. Whitten, D. S. Penneys, N. H. Williams, K. M. Neubig, L. Endara. 2006. A simple and safe method for rapid drying of plant specimens using forced-air space heaters. Selbyana. 27(1): 83–87.

Gostel, M. R., C. Kelloff, K. Wallick, AND V. A. Funk. 2016. A Workflow to preserve genome-quality tissue samples from plants in botanical gardens and arboreta. Applications in Plant Sciences 4: 1600039.

## Literature Cited

Jordon-Thaden, I. E., A. S. Chanderbali, M. A. Gitzendanner, AND D. E. Soltis. 2015. Modified CTAB and TRIzol protocols improve RNA extraction from chemically complex Embryophyta. Applications in Plant Sciences 3(5): 1400105.

